# Wideband Tuning and Deep-Tissue Spectral Detection of Indium Phosphide Nano-Laser Particles

**DOI:** 10.1101/2024.11.29.626128

**Authors:** Sangyeon Cho, Wonjoon Moon, Nicola Martino, Seok Hyun Yun

## Abstract

Laser particles (LPs) emitting narrowband spectra across wide spectral ranges are highly promising for high-multiplex optical barcoding. Here, we present LPs based on indium phosphide (InP) nanodisks, operating in the near-infrared wavelength range of 740-970 nm. Utilizing low-order whispering gallery resonance modes in size-tuned nanodisks, we achieved an ultrawide color palette with 27% bandwidth utilization and nanometer-scale linewidth. The minimum laser size was 430 nm in air and 560 nm within the cytoplasm, operating at mode order 4 or 5. We further demonstrated spectral detection of laser peaks with high signal-to-background ratios in highly-scattering media, including 1-cm-thick chicken breast tissue and blood vessels in live mice.

## 1. Introduction

Near-infrared (NIR) spectral ranges have gained increasing attention for expanding multiplexing capabilities beyond the conventional visible fluorescence spectrum and enabling deep tissue imaging.^[1–3]^ In biological tissues, optical absorption decrease significantly at wavelengths above 750 nm, and light scattering also steadily declines with increasing wavelength.^[4]^ This makes NIR light particularly advantageous for biological imaging. The NIR spectrum is typically divided into two subregions: NIR-I (750-1000 nm) and NIR-II (1000-1700 nm, also known as the short wavelength infrared or SWIR), with detection commonly achieved using silicon (Si) for NIR-I and indium-gallium-arsenide (InGaAs) photodetectors for NIR-II. Various types of NIR fluorophores have been developed, including fluorescent proteins^[5]^, single-walled carbon nanotubes^[6]^, quantum dots^[7]^, lanthanide-doped nanoparticles^[8]^, organic dyes, and polymers^[9]^. However, these conventional luminescent reagents typically exhibit emission linewidths of 50-200 nm in the NIR range, limiting their potential for applications requiring high spectral resolution.

Standalone micro- and nanoscale lasers, known as laser particles (LPs), have recently emerged as innovative optical emitters for biological applications. LPs generate much narrower linewidths (< 1 nm) than traditional fluorophores, making them ideal for applications such as large-scale spectral barcoding^[10–13]^ and spectral sensing^[14,15]^. These particles are typically fabricated from III-V semiconductor alloys in discoidal shapes, with optical resonances that can be precisely tuned by adjusting particle size. This design allowed LPs to achieve a wide emission range, spanning 30-50 nm in the far-red spectrum (650-730 nm) per semiconductor composition^[16]^ and up to 80-120 nm per alloy in the NIR II range^[10]^.

LPs with whispering gallery (WG) mode lasing can achieve wavelength scale dimensions (e.g., 1.6-2 µm for NIR-II). While these sizes are too large for molecule-specific targeting, they are well-suited for tagging cells and intracellular applications. LP emission is exceptionally bright, equivalent to the output of over 10,000 fluorophores with a submicron volume, without concentration quenching or photobleaching. This high brightness, combined with narrowband emission, ensures strong signal-to-noise performance and effective rejection of spectrally broad background noise. These unique properties make LPs highly promising for cell barcoding and sensing applications within scattering tissues, live animals, and *in vitro* environments.

Here, we present LPs made from indium phosphide (InP) that operate within the NIR-I spectral range. While InP is widely recognized for its applications in bulk and nanowire lasers^[17]^, its potential in free-standing LPs has not been explored until now. Our findings reveal that the high gain of InP enables low-order WG mode lasing, facilitating ultrawide tuning of the lasing wavelength across different mode orders. By varying particle sizes from 450 to 1000 nm, we achieve a tunable emission range from 740 to 970 nm. Leveraging from the high output intensity and background noise rejection of these LPs, we demonstrate their effective detection within cells, thick tissues, and blood streams in live mice.

## 2. Results

### 2.1 Particle fabrication and lasing characteristics

To fabricate InP nanodisks, we used a wafer that consists of InP layers and sacrificial InGaAsP layers grown on an InP substrate (**Figure** 1a, and Methods). The thickness of each InP layer was 340 nm. Initially, pillars with diameters ranging from 900 to 1100 nm were fabricated by optical lithography and reactive ion etching. Subsequent wet etching with a piranha acid solution (H_2_SO_4_:H_2_O_2_:H_2_O=1:1:10) removed the sacrificial layers, producing free-standing InP particles (Figure 1b). The particles were transferred onto a glass substrate and characterized with a home-built hyperspectral microscope. The setup employed a frequency-doubled Nd:YAG nanosecond laser (532 nm wavelength, 5 ns pulse width) and a picosecond laser (765 nm, 70 ns pulse width) for pumping, and electron-multiplying silicon cameras, time-correlated single photon counting hardware, and a diffraction grating-based spectrometer for output characterizations.

**Figure 1.**
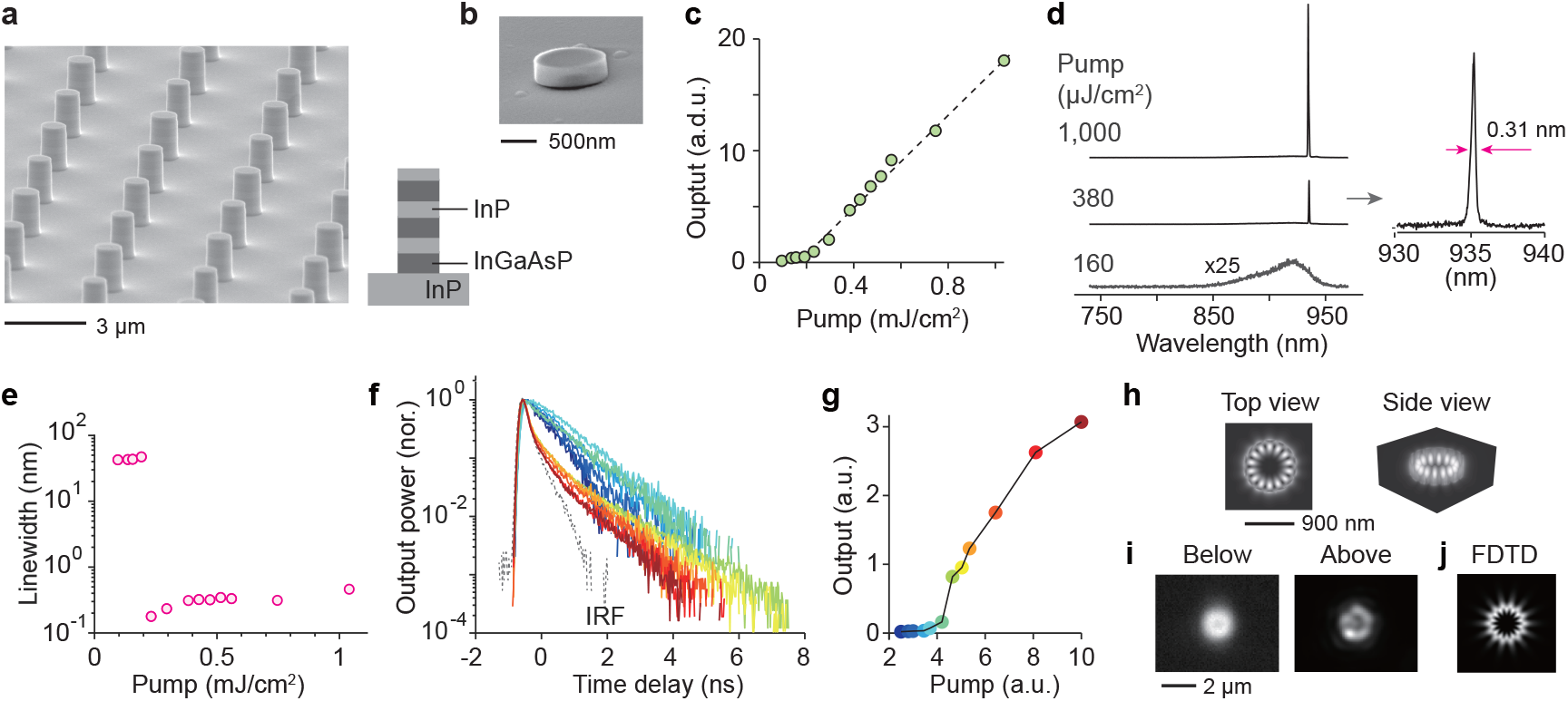
Output characteristics of InP disks. (a) Scanning electron micrograph (SEM) of fabricated micropillar arrays on a wafer. Inset: wafer design. (b) SEM of an isolated InP particle. (c) Measured light-in-light-out curve. Dashed line: theoretical fit. (d) (Left) Output spectra at different pump fluences: 160, 380, and 1000 µJ/cm^2^. (Right) A zoomed-in view of the spectrum at 380 µJ/cm^2^. (e) Measured output linewidth. (f) Transient lifetime decay curves from InP particles with a mean diameter of 900 nm. Colors correspond to different pump powers, increasing from cyan to dark red. (g) Light-in-light-out curve for the sample in (f), with the same color encoding for different pump levels. (h) FDTD-simulated electric field amplitude, |E|, of the lasing mode with an azimuthal mode order of 7 at a wavelength of 940 nm. (i) Measured far-field emission profiles below and above lasing threshold. (j) FDTD simulation for the far-field profile of the lasing mode.

The light-in-light-out curve exhibited the expected threshold behavior with a spontaneous emission β-factor of 10^−3^ (Figure 1c). Below the threshold, the photoluminescence (PL) spectrum showed a broad spontaneous emission with a 38 nm width. Upon surpassing a pump energy of 0.22 mJ/cm^2^, a sharp single-mode laser peak emerged (Figure 1d). The linewidth was 0.2 nm at the threshold Figure 1e), beyond which the linewidth increased modestly at higher pump powers. During nanosecond pumping, the decay time of the output decreased significantly above the lasing threshold (Figure 1f), when stimulated emission became dominant over spontaneous emission (Figure 1g). While the spontaneous emission decay time was approximately 1 ns, the stimulated emission decay time above the threshold was < 120 ps, limited by the instrument temporal resolution. According to finite-difference time-domain (FDTD) numerical simulation, the lasing mode expected for a disk diameter of 900 nm is the seventh-order transverse-electric WG mode (Figure 1h), with a quality factor (Q) of ∼ 360 in air. This mode has the highest Q factor among several other modes within the gain bandwidth. Experimental Q factors may be lower due to surface roughness and shape imperfections. Wide-field imaging of experimental InP particles showed ring profiles (Figure 1i), expected for the WG mode (Figure 1j).

### 2.2 Ultrawide tuning of lasing wavelength by size

We initially prepared InP microdisks with diameters of 1000 ± 60 nm (batch v). To achieve size reduction, the harvested microdisks underwent additional wet etching in an 85% wt H_3_PO_4_ solution at 70°C, with an etching rate of 17 nm/sec (34 nm/sec in diameter). By varying the etching time, we created additional batches with different mean sizes: 900 nm (batch iv), 730 nm (batch iii), 560 nm (batch ii), and 480 nm (batch i) (**Figure** 2a). The size variation within the original batch proportionally decreased with size reduction. The etching rate of InP is higher along the [001] and [010] planes compared to the [011] plane.^[18]^ Due to this anisotropic etching, the shape of the disks transformed from near-circular (batch v) to ellipsoid (batch iv) and eventually to cube-like shapes (batches i-iii).

Figure 2b presents the above-threshold spectra from a total 880 disks across the five batches. As the disk size decreased, the lasing wavelength gradually blue-shifted, reaching 740 nm for batch i. The longest observed wavelength was 970 nm in batch v. This resulted in a total tuning range of 230 nm (approximately 100 THz), corresponding to a frequency bandwidth utilization of 27%, calculated as the tuning range divided by the center frequency. This tuning range is comparable to that of bulk Ti:Sapphire lasers, which are known for their exceptionally wide tunability. The mean lasing threshold energy was 150 µJ/cm^2^ for batch v and increased to 4 mJ/cm^2^ for batch i (Figure 2c). The linewidth also broadened from ∼0.3 nm in batches v-iii (Figure 2d) to 1.5 nm in batch i. The Q factor (Q) of the lasing mode, as calculated using FDTD simulations, decreased from ∼360 in batch v to 80 in batch i (Figure 2e). Lasing involved several WG mode orders, which decreased progressively from batch v to batch i. The Mie scattering diagram (Figure 2f) illustrates the tuning curves of different modes for different batch sizes. The azimuthal mode order decreases from 7-8 in batch v to order 4 in batch i in air. In comparison, typical InGaAsP microdisks in the NIR-II range operated at the azimuthal mode order of 10-11.^[10]^

**Figure 2.**
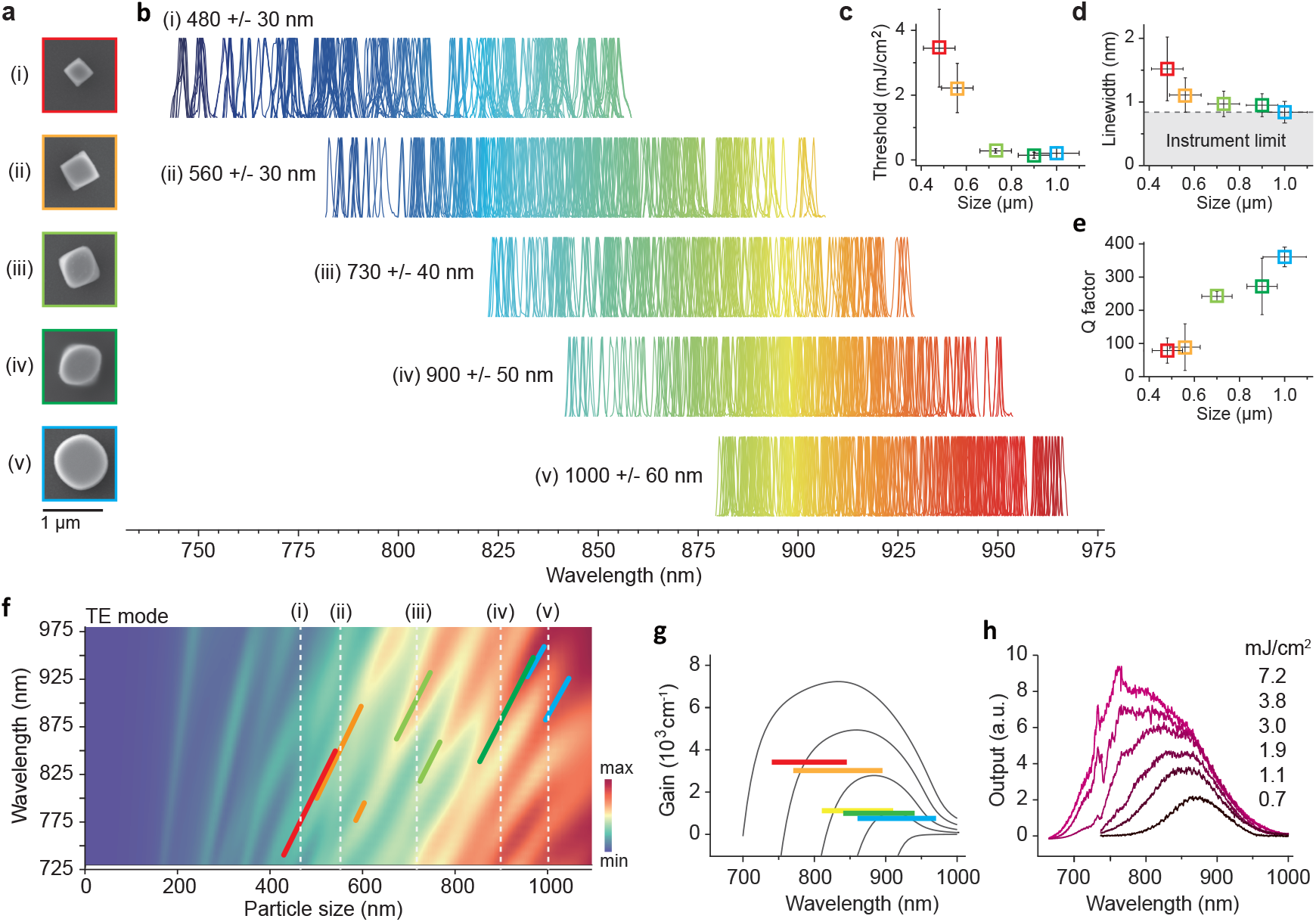
Size-dependent characteristics of InP LPs. (a) SEM images of representative particles from different batches with different sizes: (i) 480 +/−30 nm, (ii) 560 +/−30 nm, (iii) 730 +/−40 nm. (iv) 900 +/−50 nm, and (v) 1000 +/−60 nm. (b) Normalized emission spectra from numerous particles within each batch. (c) Measured lasing thresholds for each batch. (d) Measured lasing linewidths for each batch, with a spectrometer resolution of 0.8 nm. (e) Calculated Q factors of the lasing modes. Data points in panels (c)-(e) are color-coded corresponding to their batches. (f) Light scattering simulations illustrating the evolution of different transverse-electric (TE) WG modes as a function of particle size. Vertical dashed lines indicate the mean sizes of different batches. And thick color lines represent the expected tuning curves of the lasing modes within each batch. (g) Simulated semiconductor gain spectra for carrier densities of 0.1, 5, 10, 20, 40 × 10^18^ cm^−3^. Horizontal color bars indicate the loss coefficients (2πn/λQ) of the lasing modes and their spectral ranges for different batches. (h) Measured PL spectra from a non-lasing, 200 nm-sized InP particle under various pump fluences, illustrating the relationship between increasing gain coefficients and the blue-shift of the gain spectra.

For lasing to occur, the optical gain must exceed the cavity loss of the mode. The threshold gain coefficient can be approximated as 2π*n*/λQ, where *n* is the modal refractive index and λ is the free-space wavelength. For a particle size of 480 nm, the required gain coefficient is ∼ 3,400 cm^−1^, whereas a larger particle size of 1000 nm requires only 740 cm^−1^. Gain increases with pump fluences as more free electrons and holes are generated. Using a simple semiconductor theory^[19–21]^, we calculated gain coefficients as a function of wavelength for different carrier densities. These are plotted in Figure 2g for carrier densities ranging from 10^17^ to 4 × 10^19^ cm^−3^. As the pump fluence increases, free electrons and holes populate deeper into the conduction and valence bands, resulting in higher gain at shorter wavelengths. This phenomenon explains the observed blue-shift of the lasing wavelength in smaller particles. The gain profile is similar to spontaneous emission spectra. Figure 2h shows experimentally measured PL spectra from non-lasing, 200 nm-sized InP particles on a glass substrate at various pump fluences, approaching the material damage threshold of ∼8 mJ/cm^2^. Clear blue-shifts of the peak gain range are observed with increasing pump fluence, demonstrating the correlation between increasing gain and decreasing gain-peak wavelengths.

### 2.3 Intracellular InP nanolasers *in vitro*

To enhance material and wavelength stability, we coated InP particles with silica using a modified Stöber method. **Figure** 3a shows a transmission electron micrograph (TEM) of an InP particle coated with a 20-nm thick silica layer. FDTD simulations indicate the resonance wavelength of uncoated microdisks shifts by 7.3 nm when the surrounding refractive index changes by 0.1. The 20-nm silica coating reduces its sensitivity to 0.6 nm. We also produced LPs with thicker (100 nm) silica coatings to further enhance their stability against environmental changes. To facilitate cellular uptake, we further coated the silica-coated InP particles with polyethylene imine (PEI).

HeLa cancer cells were tagged with PEI-coated LPs via incubating them in a culture well for 24 hours. Figure 3b shows bright-field and PL images of a 560-nm-sized LP (from batch ii) localized within the cytoplasm under nanosecond pumping. Single-mode lasing was observed (Figure 3b), with a threshold fluence of 2.8 mJ/cm^2^ (∼ 10 pJ per pump pulse). Due to the higher refractive index of the cytoplasm (n=1.36-1.39) than air, lasing was not observed from smaller particles in batch i. The lasing wavelength showed minimal variation (< 0.1 nm) over 1 million pulses at a 10 kHz repetition rate, demonstrating excellent stability of LPs with 100 nm-thick silica coatings within cells (Figure 3c). At repetition rates below a few MHz, the pump intensity and transient heating are expected to cause minimal perturbation on cells. Figure 3d shows two cells, each containing four intracellular LPs. The presence of multiple LPs per cell enables combinatorial spectral barcoding.^[10]^ With a spectral bandwidth of 230 nm and a spectral bin of 1 nm, the potential number of unique optical barcodes generated by four randomly positioned LPs is theoretically 114 million (230 choose 4). In a CCK-8 assay, LP-tagged HeLa cells exhibited no significant difference in cell viability over 72 hours compared to untagged control cells (Figure 3e). A Live/Dead assay confirmed excellent cell viability across all particle sizes (Figure 3f).

**Figure 3.**
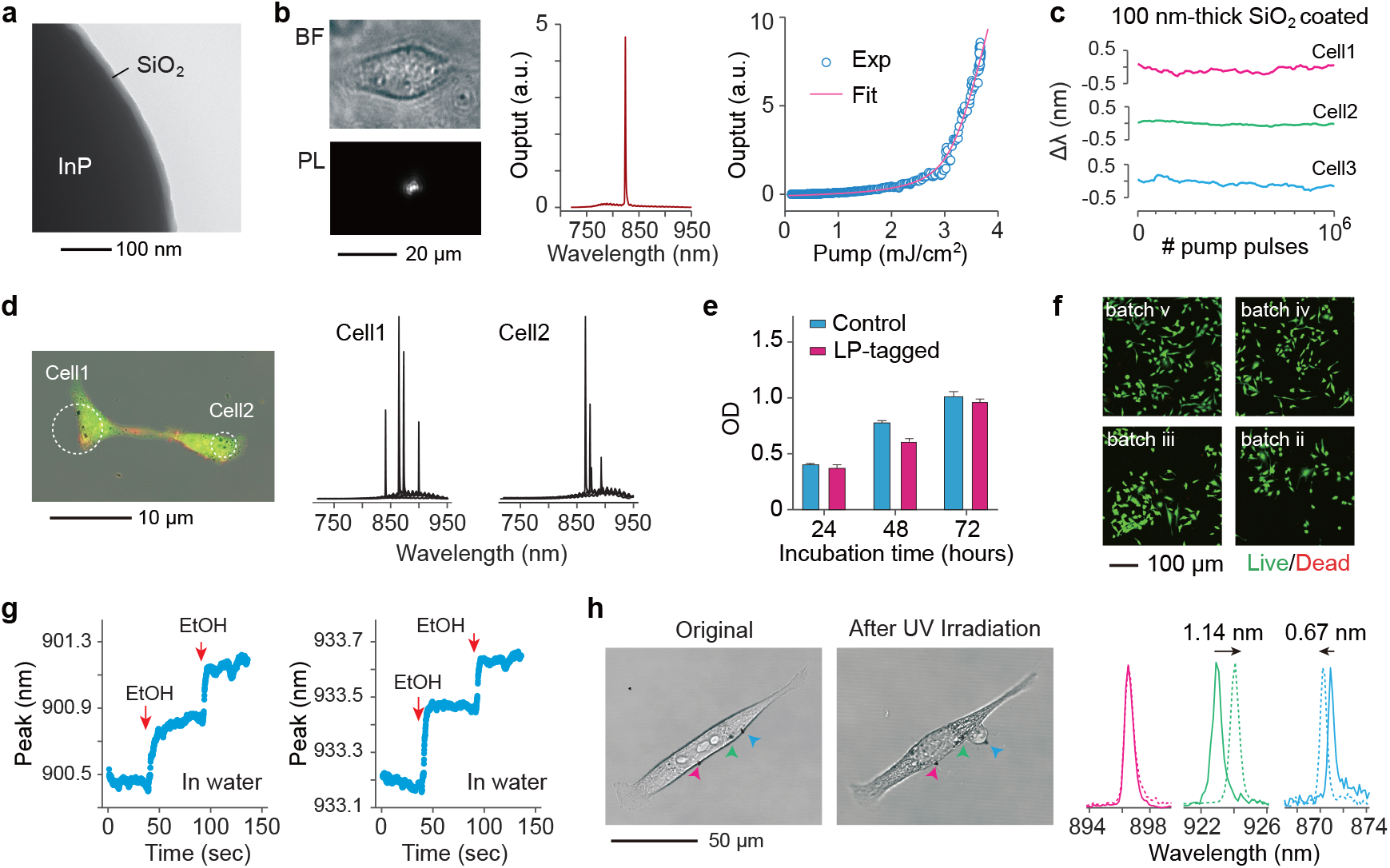
Laser characteristics within the cell. (a) TEM image of an InP disk coated with a silica layer. (b) (Left) Bright field (BF) and PL images of a 560 nm-sized LP within the cytoplasm of a live HeLa cell. (Middle) Output lasing spectrum. (Right) Light-in-light-out data. Solid curve: theoretical fit. (c) Wavelength fluctuations of three InP LPs coated with a 100 nm-thick silica layer, immersed in water during active operation with 1 million pump pulses at 10 kHz. (d) (Left) Fluorescence image of two HeLa cells (green cytoplasm dye, orange membrane dye), each tagged with four InP LPs. (Right) Output spectra from these cells. (e) CCK assay result for LP-tagged cells (blue) and un-tagged control HeLa cells (red). (f) Live (green) and dead (red) assay images of LP-tagged cells from four different batches. (g) Wavelength shifts of two InP LPs without silica coating in a reaction bath with 100 µL of water, following two sequential additions of 20 µL of ethanol (indicated by arrows). (h) (Left) BF images of a Hela cell before and after apoptosis induced by UV irradiation. Red arrow indicates an LP on the membrane, green indicates an LP near a rupture region, and cyan indicates an LP near a protruding membrane. (Right) Output spectra from these LPs before (solid curves) and after (dashed curves) the UV irradiation.

While thick silica coatings are desirable for wavelength stability, thin- or non-silica coatings offer high sensitivity to the surrounding medium for potential sensing applications. Figure 3g demonstrates a proof-of-concept experiment where two non-coated InP particles in 100 μl of water (n=1.33) on a glass-bottom dish exhibited wavelength shifts upon the sequential addition of 20 μl of ethanol (n=1.36). The measured mean shifts were of 0.28 nm for device 1, and 0.30 nm for device 2 following the first addition, and 0.22 nm for device 1 and 0.19 nm for device 2 after the second addition, similar to the theoretical shifts of 0.29 and 0.22 nm, respectively, as predicted by FDTD simulations. In another example, cells were exposed to intense ultraviolet (UV) light to induce apoptosis, and the wavelength shifts of three particles were monitored: one attached to the cell membrane and two embedded within the cytoplasm (Figure 3h). The LP outside the cell membrane showed no wavelength changes (cyan curves), while the cytoplasmic LP located near a ruptured region showed a red-shift of 1.14 nm (green curves), attributed to an increase in the surrounding refractive index by 0.016. The third LP, positioned within a cytoplasmic protrusion, exhibited a blue shift of 0.67 nm (blue curves), corresponding to a decrease in the surrounding refractive index by 0.009. These experiments highlight the potential of high-refractive-index semiconductor LPs as refractive-index sensors for both intracellular and extracellular environments, demonstrating their capability for real-time monitoring of dynamic biological processes.

### 2.4 Pumping and detection of LPs through scattering media

We investigated the excitation and detection of LPs through highly scattering media. An InP particle (from batch ii) was placed on a glass bottom dish, with translucent tape (3M Magic Tape) attached beneath the dish to simulate scattering media (**Figure** 4a). The dish was positioned on an inverted microscope and illuminated with 765 nm, 70-ps pump pulses at a repetition rate of 2.5 MHz using a 0.6-NA objective lens. The emitted light was imaged onto a silicon electron-magnification charge-coupled device (EM-CCD) camera with a 100 ms exposure time as layers of tape were sequentially added. Figure 4b shows representative CCD images. Before adding the tape layers, the LP emission form a small spot with ring interference patterns caused by the 170-µm-thick glass substrate. After the first tape layer (80 µm thick) was applied, speckle patterns began to dominate the images. Autocorrelation analysis indicated that spectral narrowing during lasing increased speckle contrast by approximately 5-fold. This trend persisted with additional tape layers, up to 17 layers (1.36 mm). The speckle pattern sizes increased with the square root of the number of tape layers (Figure 4c), characteristic of a photon diffuse regime. Correspondingly, the lasing threshold pump energy rose approximately quadratically with increasing tape thickness (Figure 4d).

**Figure 4.**
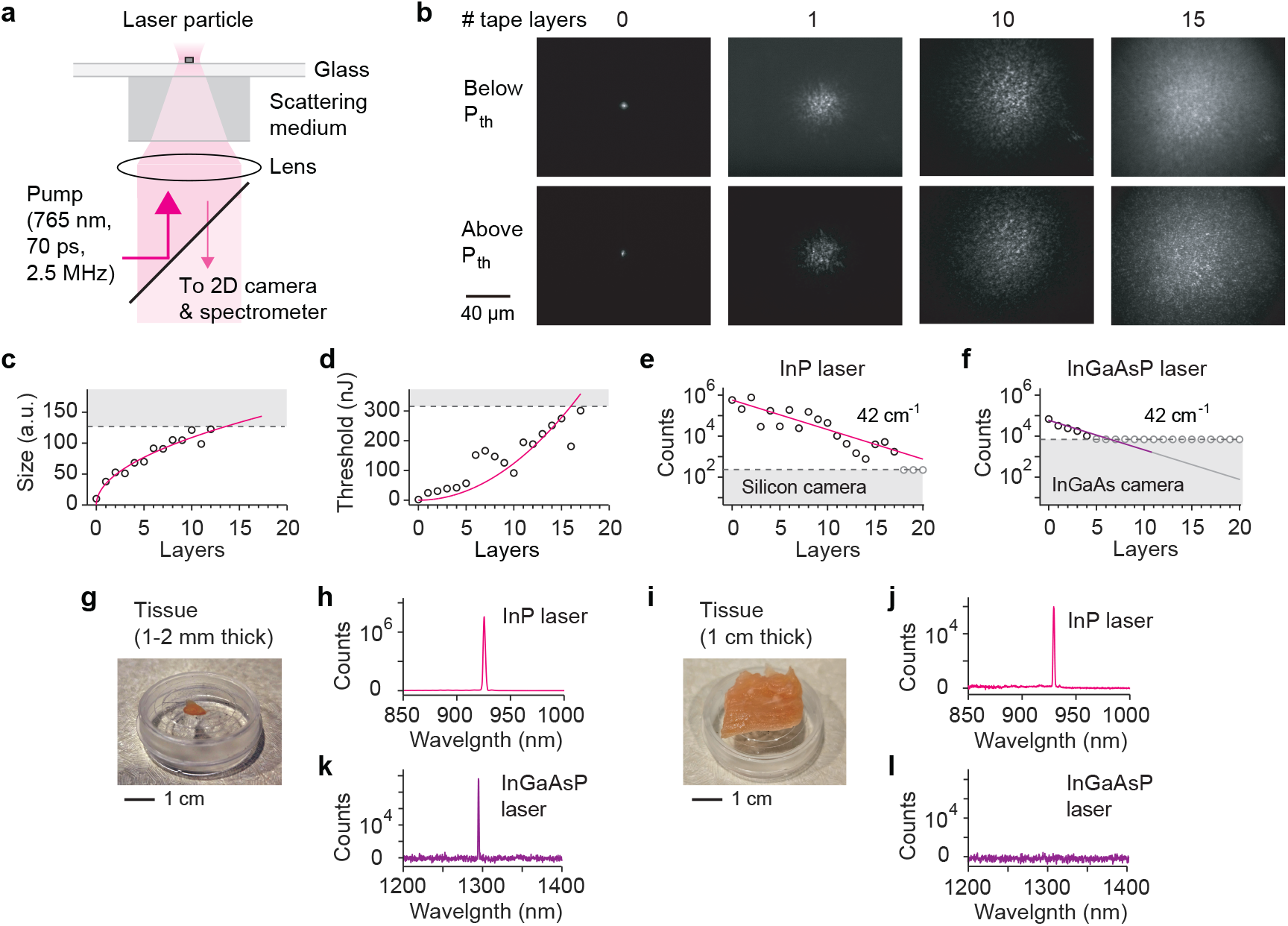
Detection of laser emission through scattering media. (a) Schematic of epi-pumping and detection of an LP through scattering media, including tape layers (b-f) and chicken breast tissues (g-l). (b) Representative widefield images of output emission from a 560-nm-sized InP laser through different numbers of tape layers, shown below and above lasing threshold. (c) Speckle pattern size as a function of tape layer thickness. The dashed line represents the measurement limit imposed by the finite size of the EM-CCD imager. (d) Threshold pump pulse energy required for detecting laser peaks as tape layers are added. The dashed line indicates the maximum energy available from the pump laser. (e) Photoelectron counts of laser peaks from an InP LP. The dashed line indicates the noise floor of the silicon EM-CCD camera in the NIR-I line-confocal spectrometer. (f) Photoelectron counts of laser peaks from an InGaAsP NIR-II LP. The dashed line indicates the noise floor of the InGaAs CCD camera in the NIR-II line-confocal spectrometer. (g) Photograph of a chicken breast tissue slice with a thickness of 1-2 mm. (h) Representative spectrum measured from an InP L through the 1-2-mm thick tissue slice. (i) Photograph of a chicken breast tissue sample with a thickness of 1 cm. (j) Representative spectrum measured from an InP LP through the 1-cm-thick tissue sample. (k) Representative spectrum measured from an InGaAsP NIR-II LP through the thinner tissue sample in (g). (l) Representative spectrum measured from InGaAsP NIR-II LPs through the thicker tissue sample in (i). No laser peaks were detected.

We recorded the photoelectron counts of laser peaks using a line-confocal spectrometer EM-CCD camera at pump energies 1.5 times the threshold values for varying numbers of tape layers. The counts exhibited an exponential decrease with a decay coefficient of 42 cm^−1^ (Figure 4e). For comparison, a similar tape experiment was conducted with a 2-µm-sized InGaAsP microdisk lasing at ∼1300 nm under 3-ns pump pulses (2 MHz). The NIR-II particle was detectable through only four layers (Figure 4f), primarily due to the lower sensitivity (∼370 photoelectrons per count, 1 ms exposure) of the InGaAs line-confocal spectrometer CCD camera used for detection.

Next, the tape was replaced with hydrated chicken breast tissues of two different thickness: 1-2 mm and 1 cm. Lasing of an InP particle was successfully achieved through both tissue samples, and narrowband emission spectra were detected with high signal-to-noise ratios (SNR) (Figures 4g-j). For the 1 cm-thick tissue, the average threshold pump power was less than 10 mW. In comparison, lasing of an InGaAsP microdisk was observed only in the thinner sample and not in the thicker one, primarily due to the lower efficiency of the InGaAs camera (Figures 4k-l).

These results highlight the advantages of narrowband lasing emission for deep-tissue detection. The sub-nanometer linewidth of the laser emission makes it easily distinguishable from broadband noise sources, such as autofluorescence, CCD electrical noise, and the spontaneous emission background of LPs. For example, using a 0.3-nm detection bandwidth captures the entire laser emission, while reducing inherently broadband fluorescence noise (e.g., with a 60-nm bandwidth) by 200-fold. This results in a 23-dB enhancement of SNR compared to non-spectral detection or luminescent particles lacking laser emission.

### 2.5 Spectral detection of Intra-tissue LP emission

To evaluate the ability to detect InP LPs within biological tissues, we performed a series of experiments. 4T1 murine breast cancer cells expressing green fluorescent protein (GFP) were tagged with PEI-coated LPs (from batch ii) at a 1:1 cell-to-LP mixing ratio. Emission spectra from 137 tagged cells on a culture dish were recorded. The tagged cells were then harvested and injected into the mammary fat pad of an anesthetized Balb/c mouse (**Figure** 5a). Confocal fluorescence microscopy (491-nm excitation) at the injection site visualized the GFP-expressing cells. Using nanosecond pumping at 532 nm, laser peaks were detected from the LPs in the injected cells. Figure 5b shows the emission spectra of three representative cells with distinct lasing wavelengths, which matched to three spectra in the dataset recorded *in vitro* prior to injection (Figure 5b, orange curves). The spectra detected from the fat pad *in vivo* exhibited slight broadening, attributed to the wider slit width used in the spectrometer to optimize the SNR.

**Figure 5.**
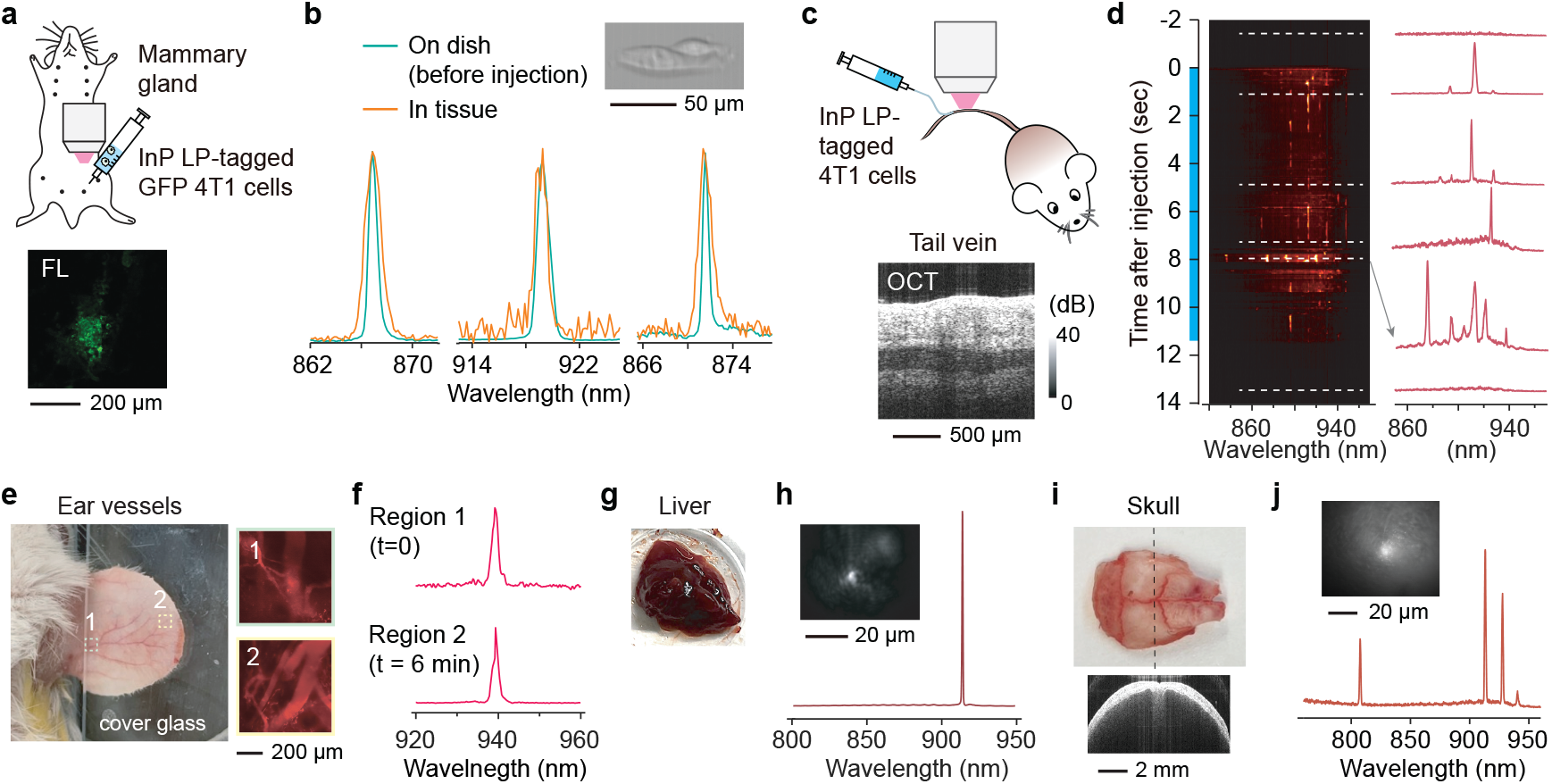
Spectral detection of intra-tissue LP-tagged cells. (a) Schematic of LP-tagged GFP-expressing 4T1 cells injected into the mammary fat pad of a mouse. Inset: confocal GFP-fluorescence image of cells at the injection site. (b) Emission spectra recorded prior to injection from three cells on a dish plate (green curves; inset: bright-field image) and matching output spectra from the same cells within the fat pad after injection (orange curves). (c) Schematic of imaging LP-tagged cells flowing in the tail vein *in vivo*. Inset: optical coherence tomography (OCT) image of the injection site, located 1 mm upstream of the imaging site. (d) Time-lapse spectral density plots recorded from the tail vein during the injection period (blue bar). Representative spectra recorded at six time points (dashed lines; 50 ms exposure each) are displayed on the right. (e) Photograph of the murine ear and fluorescence images of vasculature at regions 1 and 2. (f) Spectra recorded from an apparently identical cell in region 1 and in region 2 six minutes later. (g) Photograph of the lung collected from the mouse after euthanasia. (h) Lasing spectrum recorded from the lung tissue. Inset, wide-field PL image. (i) Photograph of the murine skull *ex vivo* alongside its OCT image. (j) Lasing spectrum from LP-tagged cells excited and detected through the skull. Inset: wide-field PL image of the specimen.

We also injected LP-tagged 4T1 cells into the tail vein of a mouse using a syringe needle (Figure 5c). An optical coherence tomography image of the injection site visualized a blood vessel approximately 200 µm in diameter located ∼ 800 μm beneath the skin surface (Figure 5c). During the slow injection of cells, a 0.45 NA objective lens was used to excite and detect LP emission 1 mm downstream, using pump pulses with 10 µJ energy at a 10 kHz repetition rate. Figure 5d shows spectral density plots acquired at 20 Hz over 16-second period (injection started at 0 s and ended at 11.3 s). Distinct individual spectra persisted for ∼ 0.2 seconds within the microscope’s field of view of 640 μm × 800 μm, suggesting an estimated flow speed of 2-4 mm/s, consistent with typical tail vein flow velocities.^[22]^ Representative emission spectra recorded at six time points (dashed lines) are shown in Figure 5d.

In a separate experiment, we focused on blood vessels in the mouse ear (Figure 5e). To visualize the ear vasculature before LP imaging, rhodamine-dextran was intravenously injected and imaged using two-photon microscopy (Figure 5e). Subsequently, LP-tagged 4T1 cells were injected into the tail vein, and 532-nm pump pulses were used to excite LPs flowing within the ear vessels. Figures 5f shows a spectrum recorded at location 1 (marked in Figure 5e) and another spectrum recorded 6 minutes later at location 2. Each spectrum persisted approximately 2 seconds within the field of view, corresponding to a flow velocity of 0.32-0.4 mm/s, consistent with typical micro-venous flow rates.^[23]^

After the ear imaging, the mouse was euthanized, and internal organs were harvested for further imaging to detect LP signals. Among the detected lasing peaks, 76% were found in the lung (Figure 5g), 18% in the liver, 3% each in the heart and spleen. Figure 5h shows a representative PL image and lasing spectrum obtained from the lung. In another experiment, lasing peaks from LP-tagged cells were excited and detected *ex vivo* through the highly scattering murine skull (Figure 5i) with high SNR (Figure 5j). These results demonstrate the capability of interrogating LPs in deep tissues and identifying their lasing peaks (spectral barcodes) from within biological tissues.

## 3. Discussion

Using high-gain InP semiconductor material, we have developed submicron-sized LPs that operate within the NIR-I range. The inherent gain properties of bulk semiconductors, which increase toward shorter wavelengths, enabled ultrawide tuning of the lasing wavelength from 740 nm to 970 nm by varying particle sizes from 1000 to 430 nm. Notably, batch-i particles in air and batch-ii particles in the cytoplasm operated at a WG mode order of 4, the lowest order ever demonstrated for photonic LPs without plasmonic effects.^[10]^ The broad spectral range and nanometer-scale linewidth position InP LPs as promising tools for large-scale barcoding applications.

The NIR-I spectral range is particularly advantageous for biological tissue applications. While the longer wavelengths of the NIR-II region benefit from reduced light scattering, the silicon-based detectors compatible with NIR-I are far more cost-effective and exhibit significantly lower electrical noise—1 to 2 orders of magnitude less—than their InGaAs-based counterparts. Compared to visible wavelengths below 700 nm, the NIR-I range of InP LP also avoids spectral overlap with most fluorescent molecules. For example, InP LPs pumped at 765 nm are compatible with most fluorophore-antibody reagents, enabling multi-parameter multi-pass flow cytometry.^[10]^

The narrowband emission from LPs allows for easy distinction from broadband emissions such as fluorescence from NIR dyes and tissue autofluorescence. This ensures that the spectral detection of laser peaks is largely immune to background noise, facilitating deep-tissue detection, as demonstrated with 1 cm-thick chicken tissues. With increasing imaging depth, the threshold pump energy rises due to pump-light diffusion. The measured threshold pump fluence for InP LPs was approximately 1 mJ/cm^2^ (or 10 pJ/µm^2^), a level well within acceptable limits for cell and tissue imaging.^[4]^ This is significantly lower than the fluences (10-100 mJ/cm^2^) typically used in conventional multiphoton microscopy and photoacoustic imaging^[24,25]^.

In conclusion, InP LPs operating in the NIR-I spectral range oeer significant potential for a variety of biological applications, including multi-pass flow cytometry, deep-tissue imaging, and single-cell tracking analysis. Their unique combination of ultrawide tunability, narrowband emission, minimal spectral overlap with widely used fluorophores, and compatibility with cost-effective low-noise silicon detection technologies make them a powerful tool for advanced biomedical research.

## 4. Experimental Section

### InP particle fabrication

Custom-designed semiconductor wafers with metal-organic chemical vapor deposition (MOCVD) epitaxial InP/InGaAsP layers on InP substrates were purchased from Seen Semiconductors. Mesa structures were fabricated on the wafers using optical lithography with circular mask patterns ranging in diameter from 900 to 1100 nm, followed by reactive ion etching. To vary the diameters of the InP layers, wafer chips were immersed in hydrochloric (HCl) or hydrophosphoric acid (H_3_PO_4_) solutions for pre-calibrated etching times. The sacrificial InGaAsP layers were subsequently removed using an acid Piranha solution (H_2_SO_4_:H_2_O_2_:H_2_O = 1:1:10), releasing the InP particles. Harvested InP particles were washed with ethanol and water and then dispersed in ethanol. For silica (SiO_2_) surface coating, a modified Stöber method was used.^[10]^ Structural verification of the silica-coated InP particles was performed using a transmission electron microscope (JEOL, JEM 1011) at a magnification of 1:250,000.

### Optical characterizations

Optical experiments were conducted using a home-built hyperspectral microscope system. Two pump sources were employed: a frequency-doubled Nd-YAG laser at 532 nm with a repetition rate of 10 kHz and a pulse duration of 2-4 ns, and an amplified frequency-doubled fiber laser emitting at 765 nm with a repetition rate of 2.5 MHz and a pulse duration of 70 ps. The system utilized either a 0.6 NA, 50x air objective lens or a 0.4 NA, 20x air objective lens. Emission from the sample, collected by the objective lens, passed through a dichroic mirror and a dichroic filter before being split into two paths. One path was directed to a silicon-based EMCCD camera (Luca, Andor) for wide-field imaging. The other path was directed to an EMCCD (Shamrock, Andor) in a spectrometer equipped with two gratings: Grating 1 (300 lines/mm, 500 nm blaze) with a 100 μm slit (resolution: 0.7-0.9 nm) and Grating 2 (1200 lines/mm, 500 nm blaze) with a 100 μm slit (resolution: 0.13 nm). The second output port of the spectrometer was coupled with an avalanche photodiode (APD) and a time-correlated single photon counting (TCSPC) system (Timeharp 260, PicoQuant) to perform transient lifetime spectroscopy. The instrument response function, characterized by using a picosecond pump laser, exhibited a time resolution of 120 ps.

### Numerical simulation

Finite difference time domain (FDTD) simulations were performed using commercial software (Lumerical). Mie scattering simulation employed a total-field scattered-field plane-wave source, while dipole simulations utilized both electric and magnetic dipoles placed inside semiconductor particles. Time-dependent electric and magnetic fields were recorded using densely positioned point-like time monitors within the semiconductor particles. Resonance frequencies (ϖ_!”#_) and the full-width-half-maximum spectral widths (Δϖ_!”#_) were used to calculate the quality factors (Q) of low-Q modes: *Q =* ϖ_!”#_ / Δϖ_!”#_. For high Q modes that did not decay completely within the simulation timeframe, Q values were determined from the slope of the electric field decay profiles. Three-dimensional near-field field patterns were obtained using an array of two-dimensional field monitors, while far-field emission patterns were calculated using a box monitor that encompassed the particle.

### Cell tagging

HeLa human cervical cancer cells and GFP-4T1 mouse mammary tumor cells were purchased from ATCC (American Type Culture Collection). HeLa cells were cultured in Dulbecco’s Modified Eagle Medium (DMEM) supplemented with 10% (v/v) fetal bovine serum (FBS) and 1% (v/v) antibiotic-antimycotic at 37 °C under 5% CO_2_. GFP-4T1 cells were cultured in Roswell Park Memorial Institute (RPMI) 1640 supplemented with 10% FBS and 1% antibiotic-antimycotic. For biocompatibility tests, HeLa cells were seeded in 96-well plates with a density of 3000 cells/well and cultured for 24 hours. InP particles with a mean diameter of 730 nm were then added to each well at a density of 6000 particles/well and cultured with the cells for 24, 48, and 72 hours. At each timepoint, cell viability was assessed using the cell counting kit (CCK8) assay by measuring absorbance (OD) at 450 nm. Cells cultured without particles were used as controls. Live/Dead assays were performed after 72 hours of culture with InP particles from different batches. For cell tagging experiments, InP particles were further coated with polyethyleneimine (PEI) on top of the silica layer to facilitate cell uptake. Hela or GFP-4T1 cells were first cultured for 24 hours and then co-cultured with PEI-coated particles for an additional 24 hours before further imaging or analysis.

### Ex vivo tissue experiments

4T1 cells were tagged with InP LPs from batch ii and cultured on a glass-bottom dish. Chicken breast tissues were sliced into 1∼2 mm or 1 cm sections and placed on top of sparsely distributed LPs (batch ii) on a glass substrate. Skull tissue was harvested from a murine carcass and attached to the bottom of a glass dish. For imaging InGaAsP NIR-II LPs, a laser-scanning confocal microscope (Olympus FV3000) was modified to incorporate a nanosecond pump laser at 1064 nm (Spectra Physics VGEN-ISP-POD, pulse duration 3 ns, repetition rate 2 MHz) and a NIR-II spectrometer equipped with an InGaAs linescan camera (Sensor Unlimited 2048). A NIR-optimized 20x, 0.45-NA objective lens (Olympus IMS LCPLN20XIR) was used for imaging.

### In vivo mouse experiments

BALB/c mice (female, 10 weeks, 20-25g) were purchased from Jackson Laboratory. Mice were anesthetized using an intraperitoneal injection of ketamine and xylazine. For fat pad imaging, the skin hair around the mammary gland was removed. GFP-4T1 cells tagged with InP LPs (batch ii) were injected approximately 2 mm away from the nipples to target the underlying fat pad, at an injection depth of ∼ 3 mm from the tissue surface. For tail vein injection, LP-tagged GFP-4T1 cells suspended in PBS (2000-5000 cells/50 µl) were injected using a cannula. Optical coherence tomography (OCT) imaging was performed using a custom-built system equipped with a swept laser with a center wavelength of 1310 nm. To visualize ear vasculature, a two-photon microscope (Olympus FV4000MPE) was used in conjunction with rhodamine-dextran (2,000,000 MW, Invitrogen), which was intravenously injected into the tail vein. All animal studies were approved by the Institutional Animal Care and Use Committee (IACUC) of Massachusetts General Brigham and conducted in accordance with National Institutes of Health guidelines (protocol 2017N000021).

## Supporting Information

Supporting Information is available from the Wiley Online Library or from the author.

## Acknowledgements

S.C., and W.M. contributed equally to this work. This study was supported by National Institutes of Health research grants (R01-EB033155, R01-EB034687). This research used the resources of the Center for Nanoscale Systems, part of Harvard University, a member of the National Nanotechnology Coordinated Infrastructure, supported by the National Science Foundation under award number 1541959.

## Data availability

Data is available upon reasonable request.

